# A deficiency screen of the 3^rd^ chromosome for dominant modifiers of the Drosophila ER integral membrane protein, Jagunal

**DOI:** 10.1101/2022.10.26.513935

**Authors:** Gerson Ascencio, Matthew A. de Cruz, Judy Abuel, Sydney Alvardo, Yuma Arriaga, Emily Conrad, Alonso Castro, Katharine Eichelberger, Laura Galvan, Grace Gundy, Jorge Alberto Inojoza Garcia, Alyssa Jimenez, Nhein Tuyet Lu, Catharine Lugar, Ronnie Marania, Tserendavaa Mendsaikhan, Jose Ortega, Natasha Nand, Nicole S. Rodrigues, Khayla Shabazz, Cynnie Tam, Emannuel Valenciano, Clive Hayzelden, Anthony S. Eritano, Blake Riggs

## Abstract

The mechanism surrounding chromosome inheritance during cell division has been well documented, however, organelle inheritance during mitosis is less understood. Recently, the Endoplasmic Reticulum (ER) has been shown to reorganize during mitosis, dividing asymmetrically in proneuronal cells prior to cell fate selection, indicating a programmed mechanism of inheritance. ER asymmetric partitioning in proneural cells relies on the highly conserved ER integral membrane protein, Jagunal (Jagn). Knockdown of Jagn in the compound *Drosophila* eye displays a pleotropic rough eye phenotype in 48% of the progeny. To identify genes involved in Jagn dependent ER partitioning pathway, we performed a dominant modifier screen of the 3^rd^ chromosome for enhancers and suppressors of this Jagn RNAi-induced rough eye phenotype. We screened through 181 deficiency lines covering the 3L and 3R chromosomes and identified 12 suppressors and 10 enhancers of the Jagn RNAi phenotype. Based on the functions of the genes covered by the deficiencies, we identified genes that displayed a suppression or enhancement of the Jagn RNAi phenotype. These include Division Abnormally Delayed (Dally), an heparan sulfate proteoglycan, the γ-secretase subunit Presenilin, and the ER resident protein Sec63. Based on our understanding of the function of these targets, there is a connection between Jagn and the Notch signaling pathway. Further studies will elucidate the role of Jagn and identified interactors within the mechanisms of ER partitioning during mitosis.

## Introduction

During cell division, it is well established that the genetic material in the form of condensed chromosomes is partitioned faithfully to the newly formed daughter cells. Lesser understood is the partitioning and inheritance of cytoplasmic material and organelles during cell division. Early models of organelle inheritance proposed a stochastic bulk inheritance of material, largely based on chance (Warren and Wickner 1996). However, recent studies have indicated a programmed pathway towards the inheritance of organelles similar to other factors necessary for cellular function and cell fate selection (Lowe and Barr 2007). Studies over the past decade have focused largely on the mitotic partitioning of the Golgi Apparatus and the Endoplasmic Reticulum (ER), outlining both their dramatic reorganization of these organelles in frame with the cell cycle and their connection with the cytoskeleton during mitosis (Wei and Seemann 2009; Yagisawa *et al*. 2013; Bergman *et al*. 2015; Smyth *et al*. 2015). Recently, the highly conserved ER transmembrane protein Jagunal (Jagn) was identified to be necessary for the proper asymmetric division of ER during mitosis in proneuronal cells, as inhibition of Jagn led to a symmetrical partitioning of the ER and defects in asymmetric division during early embryonic development (Eritano *et al*. 2017). Jagn, a highly conserved ER integral membrane protein, has been linked to protein trafficking, cell differentiation and the hematological disease, severe congenital neutropenia (VanWinkle *et al*. 2020). Furthermore, null alleles of Jagn display a lethality during early larval stages indicating a essential role during development (Lee and Cooley 2007). However, the molecular mechanism involving Jagn in these functions is poorly understood. To better understand the role of Jagn in ER partitioning and cell fate selection, we sought to identify factors that interact with Jagn using a genetic screening approach. Transcript knockdown of Jagn utilizing RNA interference (RNAi) in the *Drosophila* compound eye displays a pleotropic rough eye phenotype. To identify genetic interactors, we conducted a dominant modifier screen, using a collection of gene deficiencies along the 3^rd^ chromosome inconjunction with Jagn-RNAi and evaluated the rough eye phenotypes. Here, we found several genetic interactions with Jagn, including the glycoprotein Division Abnormally Delayed (Dally), the ER resident protein Sec63, and the γ-secretase subunit Presenilin (Psn). Based on previous studies involving these targets, there appears to be a connection between Jagn and established cell signaling pathways, indicating a regulatory connection in mitotic ER partitioning.

## Methods and Materials

### Drosophila Strains and Husbandry

Fly stocks and crosses were maintained at 25°C on standard Bloomington Drosophila Stock Center (BDSC) cornmeal medium. Third, left arm (3L) and right arm (3R) Deficiency (Df) chromosome kits were obtained from BDSC. The UAS-Jagn-RNAi was obtained from the Vienna Drosophila Resource Center (stock number, 108991 VDRC). Other fly strains used in this study can be found in Supplemental Table 1 (Table S1). Any strains used in this study are available upon request. The authors affirm that all data necessary for confirming the conclusions of the article are present within the article, figures, and tables.

### Deficiency Screening approach of the 3^rd^ chromosome

The Ey-GAL4 transgenic line was cross to UAS-Jagn RNAi and ∼47% of the resulting progeny displayed a rough eye phenotype. A collection of 181 deficiency lines covering the 3rd chromosome were screened for modification of UAS-Jagn-RNAi rough eye phenotype. The UAS-Jagn-RNAi line on the 2nd chromosome was crossed to deficiency lines on the 3^rd^ chromosome (Figure 3). These transgenic lines were then crossed to Ey-GAL4/Cyo line on the 2nd chromosome and the eye phenotypes of the progeny were scored. Over 60 flies that were heterozygous for UAS-Jagn-RNAi / Ey-GAL4 and each 3^rd^ chromosome deficiencies were then scored for either a normal eye or rough eye phenotype. Rough eye progeny that fell between 30% and 60% were considered within control range, while rough eye progeny that were above 60% were considered enhancers and rough eye progeny below 30% were considered suppressors. To confirm enhancers and supressors outside the control range (between 30 – 60%), we used a Person’s chi-squared test to determine a probability value and if there is a statistically significant difference between the expected (control) percentage and the observed (experimental) percentage.

### Target gene screening for potential interactors of Jagunal

A total of 18 genes from suppressors and enhancer 3rd chromosome regions were chosen for further analysis. Several genes covered by the identified deficiencies were eliminated based on overlap with neighboring deficiencies that showed no modification. Genes of interest were selected based on known biological roles described in Flybase.org (Gramates *et al*. 2022). The UAS-Jagn-RNAi line on the 2nd chromosome was crossed to mutant lines on the 3^rd^ chromosome (Figure 4). These transgenic lines were then crossed to Ey-GAL4/Cyo line on the 2nd chromosome. A total of 60 flies that were heterozygous for UAS-Jagn-RNAi / Ey-GAL4 and each 3^rd^ chromosome mutant were then scored for either a normal eye or rough eye phenotype.

### Preparation and imaging using Field Emission Scanning Electron Microscopy

Drosophila lines selected for imaging were placed into 1.5 ml Eppendorf tubes and fixed with 8% paraformaldehyde dissolved in PBS. Samples were kept in the fixative solution at 4°C for 24 hours on a rotating stage. The fixative was removed and the samples were washed 4 times with PBS at 4 °C for 10 minutes. To dehydrate the samples, anhydrous ETOH was added to the PBS solution at 4° C, beginning with a 50% concentration of ETOH. The concentration of ETOH was raised to 70%, 90%, 95% and 100% with the samples maintained at 4° C on a rotating stage for 24 hours. The final 100% ETOH step was repeated twice. To complete the dehydration process, the ETOH was removed from the samples in a critical point dryer (Tousimis Autosamdri-815(A). The dehydrated samples were individually mounted on 12.7 mm diameter Al stubs (Ted Pella Inc.) using silver epoxy (Ted Pella Inc.), and allowed to harden for 24 hours at room temperature. The samples were then coated with 15 nm of sputter-coated Au/Pd (Cressington 208HR) and stored under vacuum for 24 – 48 hours. Samples were then examined using a Carl Zeiss Ultra 55 Field Emission Scanning Electron Microscope (FE-SEM). Electron micrographs were recorded using an Everhart-Thornley detector at typical accelerating voltages of 1-2 keV.

## Results and Discussion

### Inhibition of Jagunal leads to a rough eye phenotype

Studies over the past 30 years have shown that the *Drosophila* compound eye is an excellent model for investigating cell signaling and cell fate determination (Freeman 1997; St Johnston 2002; Kumar 2018). In order to investigate if Jagn disruption affects eye development, we expressed the transgenic line, UAS-Jagn RNAi with the Eyeless-GAL4 (Ey-GAL4) driver (Brand and Dormand 1995). Eyeless is a transcription factor that is highly conserved and described as a master regulator of eye development (Halder *et al*. 1995). The Ey-GAL4 transgenic line has Eyeless enhancer sequences upstream of the GAL4 transgene, to induce tissue specific GAL4 expression in the compound eye. Inhibition of Jagn within the eye displayed a pleotropic phenotype with ∼52% expressing a normal eye indistinguishable from controls, while 48% displayed rough eye phenotype, with 5% producing a severe eye phenotype including several that were eyeless (Figure 1). Closer inspection of Ey-GAL4 / UAS-Jagn-RNAi (referred to as Jagn-RNAi moving forward) rough eye phenotype displayed several defects involved in cell fate selection and asymmetric division during eye development (Ready *et al*. 1976; Wolff and Ready 1991; Leshko-Lindsay and Corces 1997). Specifically, we observed defects in the size, shape, and patterning of the ommatidia when Jagn-RNAi was expressed (Figure 2A - D). There were several incidences of multiple bristles (Figure 2A, D arrowheads) and multiple sockets (Figure 2B arrows), as well as missing bristles and sockets. Additionally, there were examples where cell borders were not clearly defined (Figure 2C) indicating possible defects in polarity or cell division (Kumar 2012). These defects led us to hypothesize that Jagn may play a role in mitotic orientation (i.e., spindle positioning) and / or cell fate selection.

**Figure 1.**
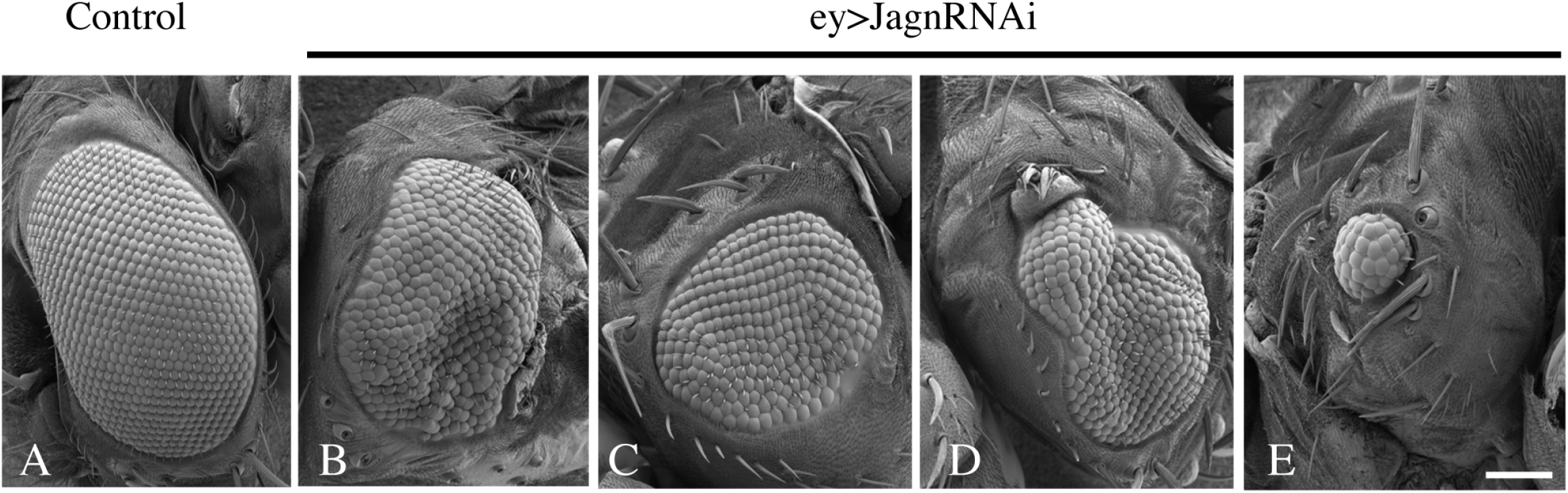
Inhibition of Jagn displays pleotropic defects in eye development. Jagn RNAi transgenic lines were crossed with the eyeless (Ey) Gal4 driver to inhibit Jagn function during eye development. Scanning Electron Micrographs (SEM) were taken of the Drosophila compound eyes and when Jagn is inhibited 52% of progeny displayed eyes similar to controls (A), while 48% displayed pleotropic eye defects (B – E). In comparison with control eyes (A), inhibition of Jagn displayed a range of defects including moderate disruption of eye shape and size (B) and (C), to more severe defects in eye development shown in (D) and (E). Scale bar ∼100 μm.

**Figure 2.**
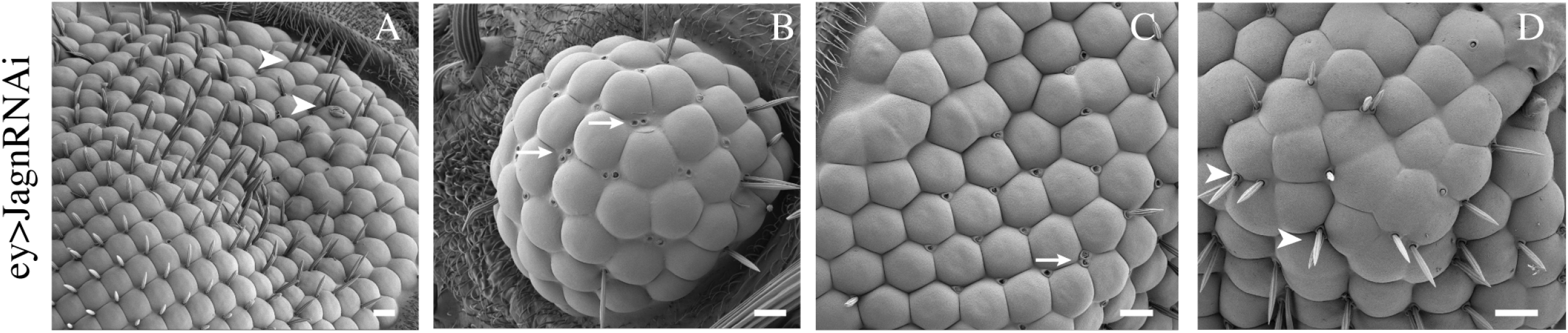
Jagn deficient compound eyes display defects in cell division and development. Examination of individual ommatidium in eyes deficient for Jagn displayed defects in cell fate selection and cell division. These include ommatidia with multiple bristles (A, D arrowsheads), or severe defects in a lack of bristles, multiple socket cells (arrows) (B), and a low number of ommatidia (B, C, D). Additionally, defects were seen in a lack of resolution of dividing ommatidia (C and D). (A) Scale Bar ∼20 μm, (B) Scale Bar ∼10 μm, (C) Scale Bar ∼10 μm, (D) Scale Bar ∼5 μm.

### Screening the 3^rd^ chromosome deficiency collection

To identify possible genes that interact with Jagn, we employed a genetic approach using a dominant modifier screen for targets that either enhance or suppress the Jagn-RNAi rough eye phenotype. This approach has been very successful in identifying components of conserved cell signaling pathways and patterning during development that traditional biochemical approaches were unable to accomplish (Banerjee *et al*. 1987; Halder *et al*. 1995). Here, we performed a crossing strategy that expressed the Jagn-RNAi transgenic line in combination with deficiency lines covering the 3^rd^ chromosome (Figure 3). These deficiencies are included in a defined kit provided by the Bloomington Drosophila Stock Center (BDSC) and cover ∼96% of the genes found on the 3^rd^ chromosome (see methods and materials). These transgenic fly populations were scored for any changes in the population expressing the Jagn-RNAi rough eye phenotype indicating either an enhancement or suppression based on the genes disrupted by the defined deficiency. Here, we screened 104 deficiency lines covering the 3^rd^ right (3R) arm of the chromosome and 77 lines covering the 3^rd^ Left (3L) arm of the chromosome. To identify modifiers of the Jagn-RNAi induced rough eye phenotype, we set a defined range of percentages of progeny that express the rough eye phenotype between 30% and 60%. Any number of progeny below 30% expressing a rough eye phenotype were considered suppressors, while any progeny above 60% were considered to be enhancers. From our screening efforts, 159 deficiencies fell within the control range (between 30-60%) and 22 were outside of the control range. To determine if these observations were significant, we performed a Person’s chi-squared analysis and determined the probability based on the expected (Jagn-RNAi control 47%) and the observed percentage of rough eye phenotype in combination with a deficiency line. Based on these criteria, we identified 12 deficiency lines that displayed a suppression of the rough eye phenotype and 10 deficiency lines that displayed an enhancement of the rough eye phenotype (Table 1).

**Figure 3.**
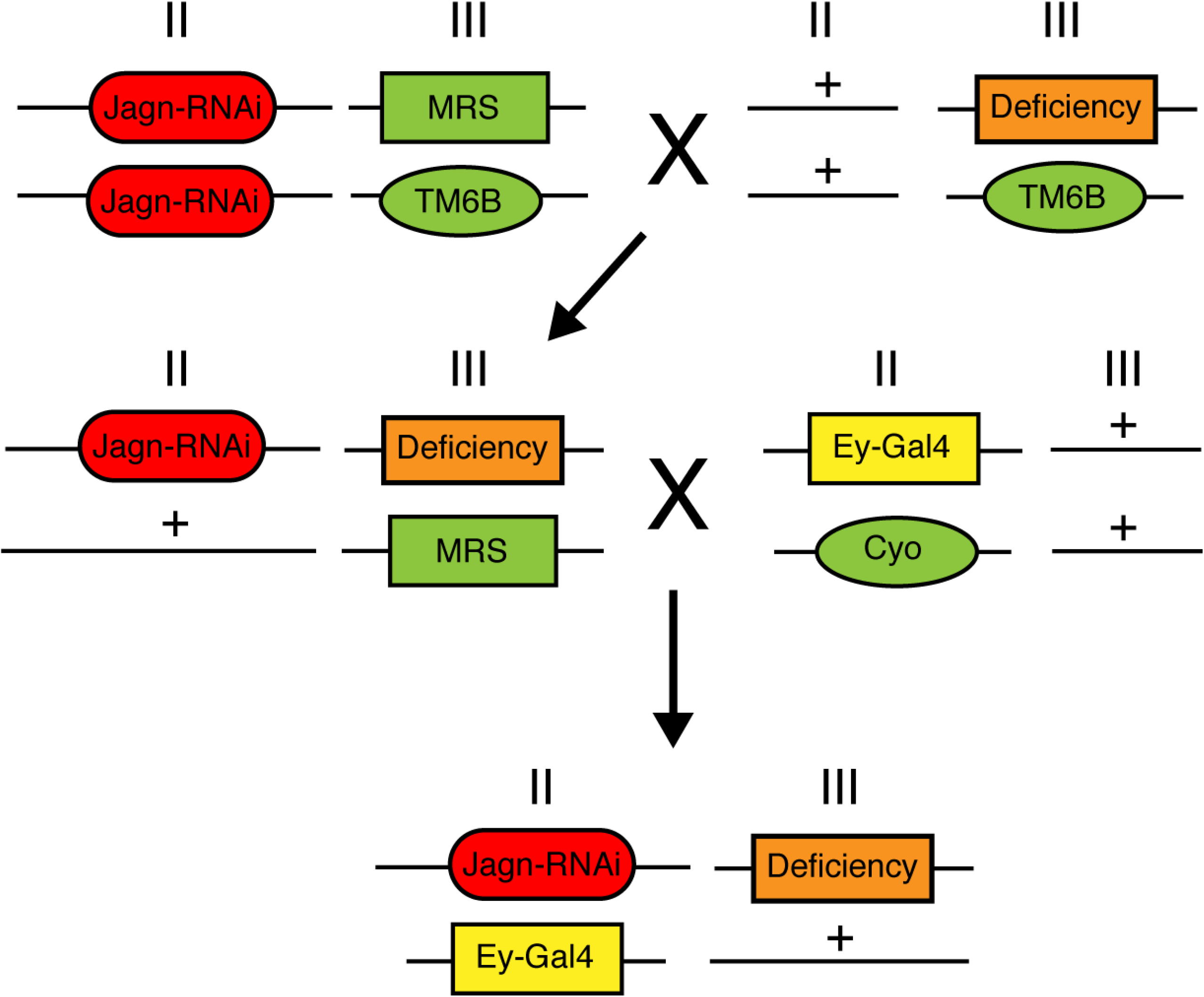
Crossing strategy for Dominant modifier screen of the 3^rd^ chromosome. In order to identify dominant modifiers of the Jagn RNAi (red) induced eye phenotype, we used the following crossing strategy to screen the collection of deficiency lines (orange) on the 3^rd^ chromosome. After two generations, a transgenic line was developed which included the Jagn RNAi line, ey-Gal4 (yellow) and a 3^rd^ chromosome deficiency. This line was screened for any enhancement or suppression of the Jagn RNAi eye defect.

**Table 1.** List of 3^rd^ chromosome deficiency lines modification of the Jagn RNAi eye phenotype. Listed are the deficiency lines covering the 3^rd^ chromosome that indicated either an enhancement or suppression of the Jagn RNAi induced rough eye phenotype. Percentages of rough eye phenotype above 60% were considered enhancers, while percentages of rough eye phenotype below 30% were considered suppressors. Control numbers of Jagn RNAi induced rough eye phenotype without a deficiency were ∼47%. N=60 eye counts for each cross. P values were determined using Person’s chi-squared analysis.

| Deficiency | Region Removed by deficiency | Effect on Jagunal RNAi Rough eye phenotype | Rough eye phenotype percentage | P-Value |
| --- | --- | --- | --- | --- |
| **3<sup>rd</sup> left chromosome** |  |  |  |  |
| **Control** | N/A | Control | 47% |  |
| **Suppressors** |  |  |  |  |
| *Df(3L)ED4177* | 61C1 to 61E2 | Suppression | 19.70% | $1.8 \times 10^{-5}$ |
| *Df(3L)BSC368* | 63F1 to 64A4 | Suppression | 25.88% | $6.5 \times 10^{-4}$ |
| *Df(3L)ED4421* | 66D12 to 67B3 | Suppression | 27.66% | 0.00158 |
| *Df(3L)BSC391* | 67B7 to 67C5 | Suppression | 29.73% | 0.00415 |
| *Df(3L)ED4470* | 68A6 to 68E1 | Suppression | 19.75% | $1.89 \times 10^{-5}$ |
| *Df(3L)ED4710* | 74D1 to 75B11 | Suppression | 26.25% | 0.000783 |
| *Df(3L)BSC775* | 75A2 to 75E4 | Suppression | 29.63% | 0.00387 |
| *Df(3L)BSC419* | 78C2 to 78D8 | Suppression | 25.88% | 0.000647 |
| **Enhancers** |  |  |  |  |
| *Df(3L)Exel6058* | 61C3 to 61C9 | Enhancement | 80.65% | $1.5 \times 10^{-5}$ |
| *Df(3L)BSC800* | 62A9 to 62A9 | Enhancement | 88.33% | $5.9 \times 10^{-8}$ |
| *Df(3L)BSC672* | 63A7 to 63B12 | Enhancement | 71.70% | 0.00215 |
| *Df(3L)ED208* | 63C1 to 63F5 | Enhancement | 91.07% | $6.3 \times 10^{-9}$ |
| *Df(3L)BSC117* | 65E9 to 65F5 | Enhancement | 83.33% | $2.4 \times 10^{-6}$ |
| *Df(3L)Exel8104* | 65F7 to 66A4 | Enhancement | 73.02% | 0.00113 |
| *Df(3L)BSC388* | 66A8 to 66B11 | Enhancement | 70.59% | 0.00359 |
| *Df(3L)BSC730* | 68F7 to 69E6 | Enhancement | 74.60% | 0.000531 |
| *Df(3L)BSC220* | 75F1 to 76A1 | Enhancement | 83.08% | $2.9 \times 10^{-10}$ |
| *Df(3L)BSC839* | 77B4 to 77C6 | Enhancement | 87.30% | $1.3 \times 10^{-7}$ |
| **3<sup>rd</sup> Right chromosome** |  |  |  |  |
| *Df(3R)BSC43* | 92F13 to 93B13 | Suppression | 20% | $2.2 \times 10^{-5}$ |
| *Df(3R)BSC650* | 90C6 to 91A2 | Suppression | 22.58% | 0.00105 |
| *Df(3R)BSC819* | 93A2 to 93B8 | Suppression | 22.7% | 0.000115 |
| *Df(3R)BSC790* | 90B6 to 90E2 | Suppression | 25.8% | 0.000624 |

The deficiency line D(3L)ED208 displayed a strong enhancement of the Jagn rough eye phenotype (*p* value = 6.3 ×10^−9^) and includes several genes involved in GPI anchor biogenesis in the ER, and oogenesis development. This is in line with previous studies on Jagn involvement in oogenesis growth (Lee and Cooley 2007). The deficiency Df(3L)BSC800 displayed an enhancement of the rough eye phenotype at 88.3% (*p* value = 5.9 × 10^−8^) of which, two genes of interest were identified, alphaCOP (aCOP) and neuronal Synaptobrevin (nSyb). aCOP is part of the COPI coat complex involved in retrograde transport and is located at the ER-Golgi intermediate complex (ERGIC) (Girod *et al*. 1999), while nSyb is a neuronal SNAP protein involved in vesicle fusion (Kidokoro 2003). Both targets are in line with a role of Jagn in ER retention and transport. This is supported by Jagn possessing a di-lysine ER retention motif (K(X)KXX) on its C-terminus. Df(3L)ED4421 showed a suppression of the rough eye phenotype (*p* value = 0.002) and has several genes of interest including Dally, an heparan sulfate proteoglycan involved in germline stem cell maintenance and Klp67A, a kinesin motor involved in chromosome segregation and mitotic spindle assembly. Df(3R)Ubx109 was also a strong suppressor and includes the genes, Arl61P which is involved in ER tubular membrane network organization and is expressed in the embryonic brain similar to Jagn. Also, the gene CG31274 is predicted to enable dynein light chain activity and is active around the centrosome and cytoplasm. The deficiency, Df(3L)BSC839 displayed a strong enhancement of the Jagn rough eye phenotype (*p* value = 1.3× 10^−7^) with 87.30% (Table 1.) containing the deleted gene, Presenilin (Psn), as well as 50 other deleted genes. In examination of two flanking deficiencies to Df(3L)BSC839, Df(3L)4858 and Df(3L)BSC797, they showed no significant changes to the rough eye phenotype, 33% and 44% respectively. This is within the control percentages, indicating no modification of the Jagn-induced rough eye phenotype by these flanking deficiencies. This led us to examine a smaller section of the Df(3L)BSC839 containing only 15 genes, with 7 being unannotated and 3 being non coding genes. This helped in narrowing down possible candidates, which included the γ-secretase subunit, Psn as an interactor with Jagn.

### Creating a targeted list of genes as potential interactors of Jagn

Based on our screening efforts described above, we identified several regions of the 3^rd^ chromosome that contain genes as possible interactors with Jagn (Table 1). Upon further examination of the genes found in these regions, there were several potential targets that we selected for further analysis. Identified genes that were found in deficiencies that showed an enhancement or suppression of the JagnRNAi induced rough eye phenotype were tested (Table 2). Identification of these genes was also aided by a previous study involving an investigation of binding partners for the human ortholog of Jagn, JAGN1 (Boztug *et al*. 2014). Studies involving human patients have linked JAGN1 to severe congenital neutropenia (SCN) and an affinity purification approach was performed identifying several factors including machinery involved in COPI vesicle formation, microtubule binding, and membrane trafficking pathways. In addition, SCN is categorized by defects in N-linked Glycosylation in primary neutrophils (Schäffer and Klein 2007). We identified several genes that are located in the deficiencies that showed a modification of the Jagn RNAi induced rough eye phenotype and sought to test these individual genes as possible interactors with Jagn. Supplemental Table 2 (Table S2) list the genes selected to test based on the biological function and mutations were genetically crossed with Jagn RNAi line and the eye phenotype was scored (Figure 4). Several of the selected targets did not show any modification of the Jagn RNAi rough eye phenotype (Table 2) including αCOP (*p* value = 0.1), Rab11 (*p* value =0.2), αTub67c (*p* value =0.02), mir-Ban (*p* value =0.2), Ar161P1 (*p* value = 0.2) and mir-282 (*p* value = 0.9). These genes were selected based on their role in the secretory pathway and neural development. In addition, there were a couple of predicted genes, CG32264 and CG739 that were of interest. CG32264 is predicted to be involved in actin binding activity and reorganization, while CG739 was a protein involved in mitochondrial translocation. Unfortunately, both did not show any modification of the rough eye phenotype. We did see a suppression of the Jagn RNAi rough eye phenotype (17%, *p* value = 4.8 × 10^−6^) with the mutation Dally (division abnormally delayed), an heparan sulfate proteoglycan (HSPs) acting as a co-receptor for growth factors and morphogens (Table 2). We also saw a suppression (28%, *p* value = 0.002) with the mutation Eip63C, a cyclin-dependent kinase that interacts with CycY and is essential for development. Mutations in Ccn (19%, *p* value = 1.9 × 10^−5^), a gene predicted to enable heparin / integrin binding activity, also displayed a strong suppression of the rough eye phenotype. Interestingly, there was an enhancement (88%, *p* value = 5.1 × 10^−8^) of the Jagn RNAi rough eye phenotype with the Sec63 mutation. Sec63 is part of a complex of proteins that are involved in the translocation of mRNAs into the ER lumen (Linxweiler *et al*. 2017; Jung and Kim 2021) and interacts with ER chaperone protein, BiP. However, the role of Sec63 in ER translocation is still poorly understood and substrates of Sec63 still remain to be identified.

**Table 2.** Modification of Jagn RNAi rough eye phenotype with selected genes. Mutations in selected genes were examined for a modification of the Jagn RNAi induced rough eye phenotype. Psn alleles [143] and [C4] and Sec63 displayed an enhancement of the rough eye phenotype, while Dally, Ccn, dar1, and Eip63E showed a suppression. Control numbers of Jagn RNAi induced rough eye phenotype without including a mutant allele were ∼47%. N=60 eye counts for each cross. P values were determined using Person’s chi-squared analysis.

| Genes | Effect on Jagunal RNAi<br>Rough eye phenotype | Rough eye phenotype<br>percentage | P-Value |
| --- | --- | --- | --- |
| Psn [143] | enhancement | 71.7% | 0.00218 |
| Psn [C4] | enhancement | 75.4% | 0.000326 |
| Eip63E | suppression | 28.36% | 0.00221 |
| dar1 | suppression | 27.87% | 0.00175 |
| Sec63 | enhancement | 88.52% | $5.1 \times 10^{-8}$ |
| Dally | suppression | 17.65% | $4.8 \times 10^{-6}$ |
| Ccn | suppression | 19.74% | $1.9 \times 10^{-5}$ |
| α-Tubulin 67C | — | 34.33% | 0.027 |
| Klp67A | — | 38.33% | 0.099 |
| α-COP | — | 38.71% | 0.110 |
| miR-ban | — | 59.34% | 0.187 |
| Sec15 | — | 44.62% | 0.447 |
| Rab11 | — | 40.30% | 0.170 |
| Arl6IP1 | — | 41.67% | 0.239 |
| CG7394 | — | 43.55% | 0.362 |
| Girdin | — | 45.16% | 0.494 |
| miR-282 | — | 49.12% | 0.901 |

**Figure 4.**
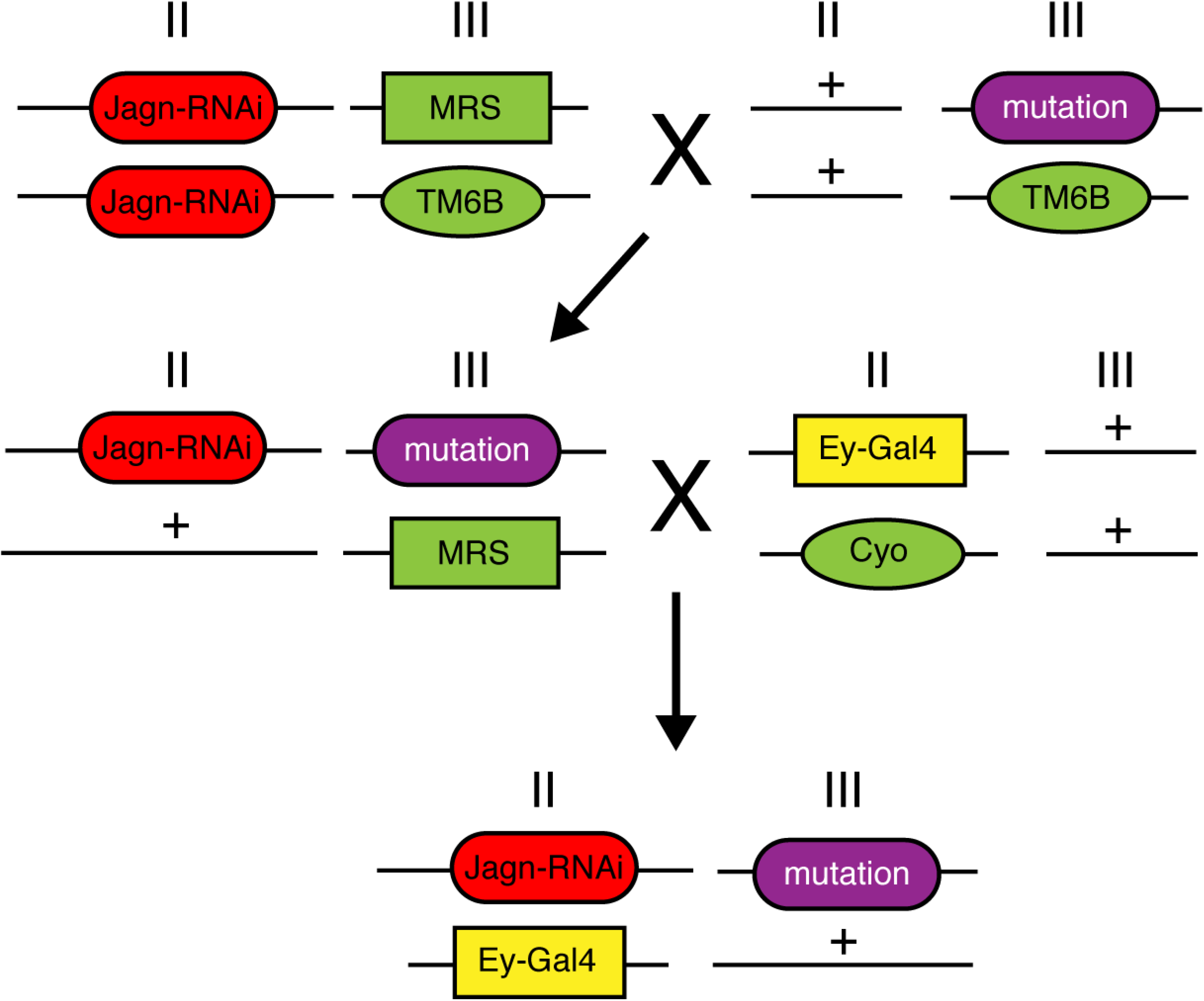
Crossing strategy of selected genes involved in modification of the Jagn RNAi induced eye phenotype. To examine specific genes involved in the modification of the Jagn RNAi (red) induced eye phenotype, we used the following crossing strategy to screen the mutant alleles of (purple) on the 3^rd^ chromosome. After two generations, a transgenic line was developed which included the Jagn RNAi line, ey-Gal4 (yellow) and the selected mutant allele. This line was screened for any enhancement or suppression of the Jagn RNAi eye defect.

Based on our examination of Df(3L)BSC839 and identification of the region including Psn, we sought to investigate if Psn displays a modification of the Jagn-RNAi induced rough eye phenotype. Support for this interaction stems from the above mentioned study involving human JAGN1 and its interaction with the protein, Adipocyte plasma membrane-associated protein (APMAP), an inhibitor of amyloid-beta (Aβ) aggregates (Boztug *et al*. 2014). Additionally, a recent study also showed that APMAP interacts with γ-secretase and is involve in modulating its activity (Mosser *et al*. 2015). Here, we crossed mutations in Psn, the Drosophila ortholog of γ-secretase (Struhl and Greenwald 1999), with the Jagn RNAi induced rough eye phenotype and examine the progeny for a modification of the eye phenotype (Table 2). We saw a strong enhancement of the rough eye phenotype (75.4%, *p* value = 0.0003) in the presence of the Psn mutation, indicating a genetic interaction between Psn and Jagn.

Overall, our screening efforts have identified several interactors with Jagn including Dally, Sec63, Eip63C, Ccn, and Psn. While this is not an exhaustive list of interactors along the 3^rd^ chromosome, these modifiers do shed some light onto the role of Jagn and indicate a possible mechanism involved the signaling and distribution of ER membrane in cells as they adopt their cell fate. Of particular interest is the HSP protein, Dally. Dally was initially identified as an integral membrane proteoglycan required for cell division during patterning formation in development (Nakato *et al*. 1995) and studies connected Dally function to Frizzled function in transducing wg signaling (Lin and Perrimon 1999). However, over the past several years, studies have also liked Dally function to other signaling pathways including TGF-beta and JAK/STAT signaling. Recently, Dally also has been linked to Notch signaling involving maintenance of the germline stem cell niche (Zhao *et al*. 2020). This indicates that Dally plays a role in several signaling pathways and is upstream of Notch signaling involving stem cell development.

Further support involving a connection between Jagn and the Notch signaling pathway, is the identification of the γ-secretase Drosophila ortholog Psn as a genetic interactor with Jagn. There have been several studies demonstrating that γ-secretase is responsible for cleavage and processing of the Notch receptor for transcription of genes involving the generation and differentiation of neuronal cells (Roncarati *et al*. 2002; Sorensen and Conner 2010). In addition, γ-secretase has also been linked to disorders including Alzheimers disease and several types of cancer (Miele *et al*. 2006; De Strooper *et al*. 2012).

Future studies of Jagn function in proneuronal cells and its role in cell fate selection will focus on the connection to the Notch signaling pathway and the generation of cell diversity in the central nervous system.

## Acknowledgements

We thank the Electron Microscopy Facility (EMF) and the Cell and Molecular Imaging Center (CMIC) at SFSU, and Dean Carmen Domingo for the provision of research facilities. We gratefully acknowledge the assistance of Dr. Annette Chan and Diana Mars. We would also like to thank the leadership at the Student Enrichment Office (SEO) at SFSU for continuing to support student engagement in research. We would like to thank San Francisco State University and the support that they have shown to our undocumented and Deferred Action for Childhood Arrivals (DACA) student population. It is through these efforts that the research in this manuscript was realized.

## Supplemental Figures and Tables

**Supplemental Table 1. List of Drosophila stocks used**. In addition to Deficiency (DF) collection covering the 3^rd^ chromosome, there were several other stocks used including additional deficiencies listed that assisted in identifying gene targets due to their overlay with deficiencies found in the kit.

**Supplemental Table 2. List of interested genes from deficiency modifiers**. Several deficiencies displayed either an enhancement or suppression of the Jagn RNAi induced rough eye phenotype. Examination of genes covered by the deficiencies produced several potential genes that interact with Jagn. Genes were identified based on known biological function provided by Flybase and areas that did not overlap with deficiencies that did not display a modification.

